# GITR and TIGIT immunotherapy provokes divergent multi-cellular responses in the tumor microenvironment of gastrointestinal cancers

**DOI:** 10.1101/2023.03.13.532299

**Authors:** Anuja Sathe, Carlos Ayala, Xiangqi Bai, Susan M. Grimes, Byrne Lee, Cindy Kin, Andrew Shelton, George Poultsides, Hanlee P. Ji

## Abstract

Understanding the cellular mechanisms of novel immunotherapy agents in the human tumor microenvironment (**TME**) is critical to their clinical success. We examined GITR and TIGIT immunotherapy in gastric and colon cancer patients using *ex vivo* slice tumor slice cultures derived from cancer surgical resections. This primary culture system maintains the original TME in a near-native state. We applied paired single-cell RNA and TCR sequencing to identify cell type specific transcriptional reprogramming. The GITR agonist was limited to increasing effector gene expression only in cytotoxic CD8 T cells. The TIGIT antagonist increased TCR signaling and activated both cytotoxic and dysfunctional CD8 T cells, including clonotypes indicative of potential tumor antigen reactivity. The TIGIT antagonist also activated T follicular helper-like cells and dendritic cells, and reduced markers of immunosuppression in regulatory T cells. Overall, we identified cellular mechanisms of action of these two immunotherapy targets in the patients’ TME.

---

The success of checkpoint blockade for cancer immunotherapy has spurred on the development of new immune-related therapeutic targets. To understand the mechanistic effects of these immunotherapy agents in cancer, one must evaluate their impact on the diverse cell types that are present within the tumor microenvironment (**TME**). Cell culture systems of T cell exhaustion and mouse cancer models are commonly used to evaluate immunotherapy agents. However, neither of these experimental approaches replicate the cellular diversity found in the native TME within patients’ malignancies (1).

In patient tumors, the native TME has a wide array of cell types including T cells, fibroblasts, macrophages, etc. Moreover, many cell types within the native TME have specific functional phenotypes that are a challenge to replicate within vitro systems. For example, TME-based CD8 T cells exhibit naïve, cytotoxic or exhausted functional phenotypes (2). These cellular phenotypic ‘states’ have a profound impact on response to an immuno-perturbation. Thus, there are significant advantages for using experimental methods that fully represent the TME cellular complexity, identify the different cell types and determine their functional states. For example, a recent study assessed the cellular effects of PD-1 blockade on *ex vivo* fragment cultures of patient tumor specimens (3). Early *ex vivo* cytokine and chemokine responses at 48 hours correlated with clinical response. Thus, using primary tissue cultures has utility for evaluating cellular responses in the TME to understand immunotherapy effects.

A method for preserving cellular composition of cancers involves tumor slice cultures (**TSCs**) (4). Tumors originating from surgical resections are rapidly processed into thin slices and then placed in culture media. The tissue sections’ thickness is in the range of several hundred microns which enables rapid diffusion of media, oxygen and other molecules. This primary tissue culture approach has been well-established in preserving the cellular TME of original tumor (4–7). While TSCs have been used to evaluate the effects of chemotherapy agents in primary tumors specimens (8–10), only a limited number of recent studies have leveraged them to determine the consequences of immunotherapies such as anti-PD-1, anti-TIM-3, anti-IL-10 and CAR-T cells (11, 12).

There are additional challenges for evaluating the impact of targeting specific immune blockade molecules. Conventional experimental methods do not provide the resolution to identify the complex features of individual TME cells. Many studies use fluorescent antibody staining approaches to identify specific cells, either through flow cytometry or microscopy. However, these methods capture a limited number of pre-defined molecular features among the affected cells. Another experimental approach involves using conventional RNA-seq to identify gene expression changes in the TME. However, standard RNA-seq requires processing the tissues in bulk and lacks the discrimination of assigning gene expression to individual cell types present in the TME. More recently, single-cell RNA sequencing (**scRNA-seq**) has provided an unbiased assessment of individual cell’s transcriptional changes. Single cell gene expression defines specific cell types or functional states. Single cell genomic methods have provided valuable information about how specific cell types respond to PD-1 blockade using longitudinal pre and on treatment patient biopsies (13, 14).

To address the challenges of studying the effects of candidate immunotherapies in the native TME, we used an integrative approach. It combines the TSC experimental model with single-cell genomics. We determined how specific antibodies targeting immune checkpoints or co-stimulatory molecules altered the immune and other cell types present in the native TME from gastrointestinal cancers. To evaluate the cellular effects, we used single cell RNA sequencing (**scRNA-seq**) and single-cell TCR sequencing (**scTCR-seq**). Across a series of TSCs derived from colorectal and gastric carcinomas, we determined the cellular response of specific immune perturbations, namely antibodies targeting specific checkpoints or co-stimulatory molecules. Single cell gene expression provided a readout to determine how specific TME cell subpopulations were affected by these perturbations.

We tested two antibodies targeting GITR and TIGIT. Antibodies targeting GITR and TIGIT are both being actively evaluated in various clinical trials for cancer (20). GITR is a co-stimulatory T cell receptor (15). TIGIT is a co-inhibitory receptor, which binds with ligands from the PVR/NECTIN family and reduces the costimulatory function of the CD226 receptor (16). Previously, we identified both these targets in a single cell genomic analysis of gastric cancers’ TME (**GC**) (17). These targets were over-expressed in both exhausted CD8 T cells and regulatory T cells (**Tregs**) in the TME but not in paired normal gastric tissue. Similar findings have been reported in colorectal cancer (**CRC**) (18) and several solid tumors (19).

We evaluated GITR and TIGIT target expression in a series of colorectal and gastric cancers using multiplex immunofluorescence (**mIF**). We also confirmed the expression of *GITR* and *TIGIT* across a scRNA-seq dataset from 217 patients that included colorectal cancers as well as 12 other tumor types (21). We also confirmed the expression of these targets from our previously published GC dataset (17).

We determined that GITR agonist antibody had only a limited cellular effect which was primarily restricted to cytotoxic effector CD8 T cells. In contrast, when we tested a TIGIT antagonistic antibody on TSCs from the same set of cancers, we observed increased TCR signaling and activation in both cytotoxic and dysfunctional CD8 T cells, including in expanded clonotypes. Moreover, using the TIGIT antagonist antibody, we observed activated follicular helper-like (**TFh-like**) cells and a reduction in the immunosuppressive phenotype of Tregs and dendritic cells (**DCs**). These results demonstrated how single cell genomics combined with TSCs can be applied to primary gastrointestinal cancers to identify the heterogenous cellular responses to GITR stimulation and TIGIT inhibition.

## RESULTS

### Experimental approach and study design

We obtained ten surgical resections of CRCs or GCs from seven different patients (**Table 1**). The tissue samples included seven resections of primary CRC from seven individual patients. From one patient with GC, we obtained three independent resections – one from the primary tumor and two from metastases to the peritoneum which is an organ that lines the abdominal cavity.

**Table 1:** Study samples.

|  |  | Experimental conditions |  |  |  |  |
| --- | --- | --- | --- | --- | --- | --- |
| Sample ID | Tumor site | T0 | ctrl | PMAIono | GITR | TIGIT |
| CRC-1 | primary CRC | + | + | + | + | - |
| CRC-2 | primary CRC | + | + | + | + | - |
| CRC-3 | primary CRC | + | + | + | + | - |
| CRC-4 | primary CRC | + | + | + | - | + |
| CRC-5 | primary CRC | + | + | + | + | + |
| CRC-6 | primary CRC | + | + | + | + | - |
| CRC-7 | primary CRC | + | + | + | + | + |
| GC1-1 | primary GC | + | + | + | + | - |
| GC-1-2 | metastatic GC | + | + | - | + | + |
| GC-1-3 | metastatic GC | + | + | - | + | + |

CRC: colorectal cancer, GC: gastric cancer, T0: original surgical resection, ctrl: control,
PMAIono: PMA Ionomycin

The samples underwent rapid processing following surgical resection (**Methods**). From each tumor, we split the tissue, using one portion to generate single cells suspensions and scRNA-seq and scTCR libraries for sequencing (**Fig. 1A, Methods**). The tissues that were processed immediately into single cell libraries provide a baseline of the cellular composition of the tumor. We refer to this baseline as the T0 time point. The other portion were used for TSC culturing. These results were used for determining cellular changes that may occur during the TSC culturing.

**Figure 1.**
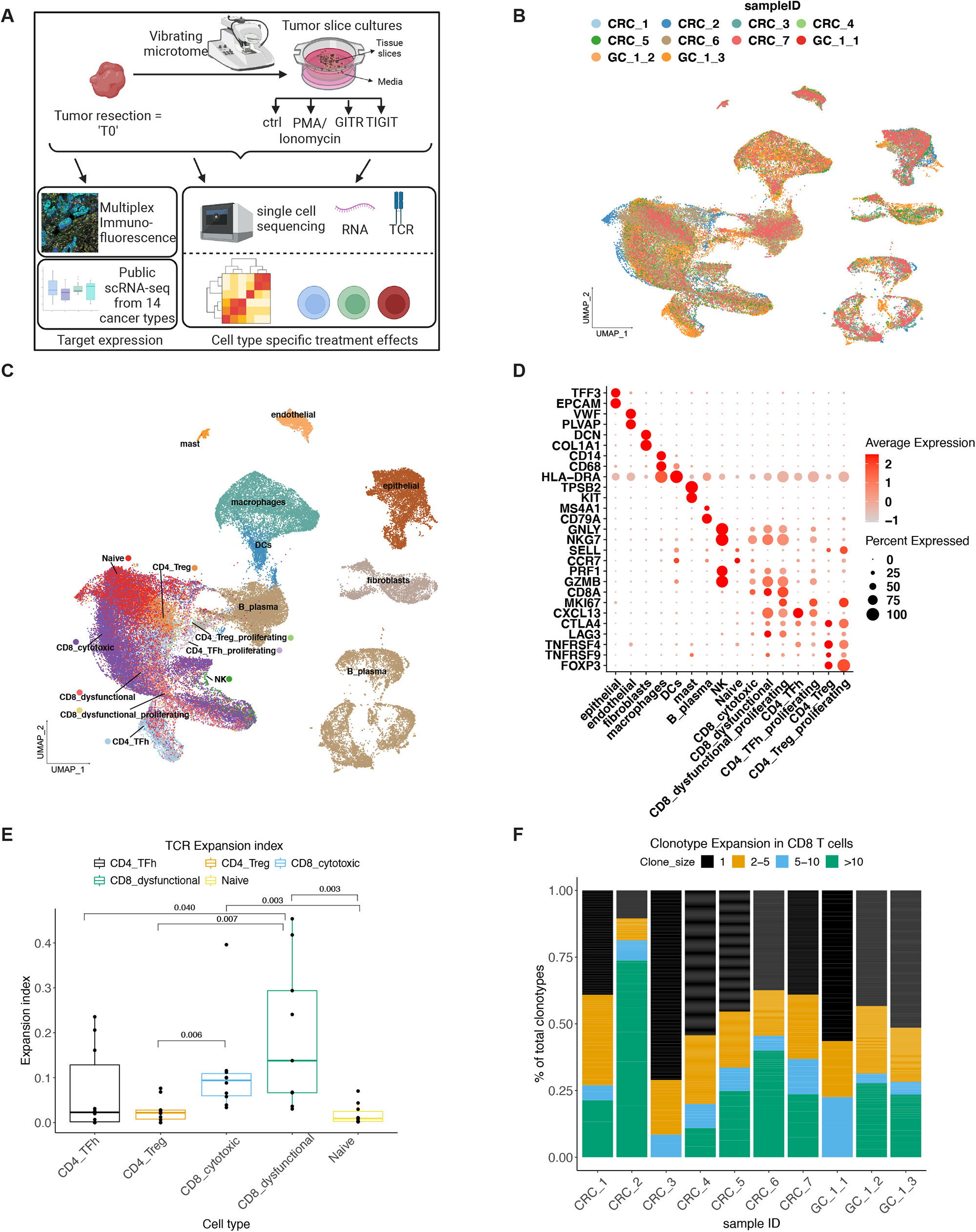
(A) Schematic representation of study design. (B-C) UMAP representation of dimensionally reduced data following batch-corrected graph-based clustering of all datasets colored by (B) samples and (C) cell type. (D) Dot plot depicting average expression levels of specific lineage-based marker genes together with the percentage of cells expressing the marker. (E) TCR expansion index for respective cell types. *p* from pairwise Wilcoxon test with Benjamini-Hochberg correction. (F) Frequencies of clonotypes in CD8 T cells from respective patients.

For experimental testing of each cancer’s TME, we generated *ex vivo* tumor slice-cultures (**TSCs**) from the resections. There were four different conditions. The TSCs were treated with either i) isotype control antibody (‘**ctrl’**), ii) T-cell activator PMA/Ionomycin (‘**PMAIono’**), iii) GITR agonist antibody (‘**GITR**’) or iv) TIGIT antagonist antibody (‘**TIGIT**’). After 24 hours of treatment, the cells were harvested and then processed for scRNA-seq and scTCR-seq. The number of experimental conditions tested per sample depended on the available size of each resection. All ten samples had adequate tissues for a baseline T0 and control samples for scRNA-seq. We conducted scRNA-seq on PMA/Ionomycin treatment from eight samples, GITR agonist treatment from nine samples and TIGIT antagonist treatment from five samples (**Table 1**). Quality control measures including filtering cells for mitochondrial genes indicative of cell death (22) and doublet identification (23). Following filtering, our final analysis included a total of 236,483 single cells with an average of 5,630 cells per sample (**Supplemental Table 1**).

### Baseline immune cell characteristics of the TME from primary gastrointestinal tumors

We determined the baseline cellular composition of the T0 samples (**Fig. 1B**). Batch effects were reduced using the Harmony algorithm (24). Specific cell type clusters were composed of different samples, indicating the elimination of batch effects (**Fig. 1C**). Using the scRNA-seq data, we made cell type assignments based on canonical marker genes (**Methods**). Overall, we identified tumor epithelium (*EPCAM, TFF3*), macrophages (*CD68, CD14*), DCs (*HLA-DRA*), mast cells (*KIT, TPSB2*), fibroblasts (*COL1A1, DCN*), endothelial (*VWF, PLVAP*) and B/plasma (*MS4A1, CD79A*), T and NK cells (**Fig. 1C, 1D**).

We characterized the T and NK functional cell states using a method called cell reference mapping. This method uses an established reference from a pan-cancer tumor immune cell atlas (21) and the SingleR algorithm (25). Each cell’s gene expression is matched to a given reference cell type. This approach provides an unbiased identification of cell subtypes without applying cell clustering methods. For these results, we denote the cell type and functional state by listing prominent examples among the associated gene expression markers. We identified different T cell subtypes including CD4/CD8 naïve cells (*CCR7, SELL, LEF1, TCF7*) (**Fig. 1C, 1D**). Among the CD8 T cells, we identified cytotoxic CD8 expressing effector cytokines (*GZMK, GZMA, PRF1, NKG7*) with low expression of immune checkpoints. Dysfunctional CD8 T cells (*LAG3, TIGIT, PDCD1, HAVCR2, CTLA4, CXCL13*) were also observed. These exhausted CD8 T cells have increased immune checkpoint expression (2). Dysfunctional CD8 T cells had a subset of proliferating cells noted by expression of the marker gene *MKI67*. Proliferating dysfunctional CD8 T cells have been linked to early dysfunction in a clonal tumor-reactive population (26).

We identified CD4 TFh-like (*CXCL13*) and regulatory T (**Treg**) cells (*FOXP3, IL2RA*). TFh-like cells have been linked to anti-tumor immunity by promoting CD8 and B cell activity (2, 27). In contrast, Treg cells are immunosuppressive and limit anti-tumor activity through specific effects on cytotoxic CD8 T cells, dendritic cells and macrophages (28). As corroborated by other studies (26), we observed proliferative subsets among the TFh-like and Treg cells. These proliferative subsets may reflect a TME response to local tumor antigens (2). These cell types were identified across all patients in varying proportions (**Supplemental Fig. 1A**). In summary, across all tissue samples, the TME in the baseline T0 resections contained diverse functional T cell states with anti-tumor (cytotoxic CD8, dysfunctional CD8, TFh-like) and immunosuppressive (Treg) properties.

### Baseline T cell receptor clonality in the primary tumor TME

To assess clonality of the T cells in the cancer’s TME at the baseline state, we performed scTCR-seq on the baseline T0 samples. We identified TCR chains from an average of 43% of the T cells with matching single cell gene expression (range 30% – 78%). Next, we determined whether there was evidence of TCR clonotype being highly represented within a given sample (18). This overrepresentation is termed as being “an expansion” for a given T cell clonotype. Moreover, one can assign specific clonotypes to different cell states (i.e., Tregs, Tfh, etc.).

To conduct this analysis, the frequency of individual clonotypes was calculated using the Shannon entropy score – this metric quantifies T cell clonotype expansion with a value range of 0 to 1, with 1 indicating high clonality. Cytotoxic and dysfunctional CD8 T cells showed significantly high expansion index of TCR clones (**Fig. 1E**). High clonality and expansion may be an indicator of tumor-antigen driven expansion in the TME (18). Next, we examined the frequency distribution of CD8 T cell clonotypes across samples (**Fig. 1F**). Single cell clonotypes represented the majority of TCRs, indicating a lack of expansion among these cells. Across the samples, between 29 to 90% of the total clonotypes were detected in more than one cell, indicating expansion. This analysis identified TCR sequences of expanded clones that may be potentially tumor-reactive in each sample at baseline. Overall, we identified TCR sequences that belonged to expanded clonotypes in infiltrating CD8 T cells in the baseline tumors.

### GITR and TIGIT gene and protein expression in the baseline TME

We evaluated the gene expression of the immunotherapy targets *TNFRSF18* (encoding protein GITR) and *TIGIT* among gastrointestinal cancers at the baseline state. In both CRCs and GC tumors, the dysfunctional CD8, TFh-like and Treg cells had the highest levels *TNFRSF18* expression (**Fig. 2A, 2B**). The complementary ligand, *TNFSF18,* encoding the protein GITRL, was expressed by fibroblasts, DCs and macrophages in CRC. *TIGIT* expression was highest in cytotoxic CD8, dysfunctional CD8, TFh-like and Treg cells. The genes *PVR* and *NECTIN2*, which encode for TIGIT ligands, were expressed by tumor epithelial, endothelial, fibroblasts, macrophages and DCs in the TME. These expression patterns are along the lines of other reports (29, 30). Overall, this result indicated that among all samples, the TME cells expressed genes required for GITR and TIGIT receptor-ligand signaling.

**Figure 2.**
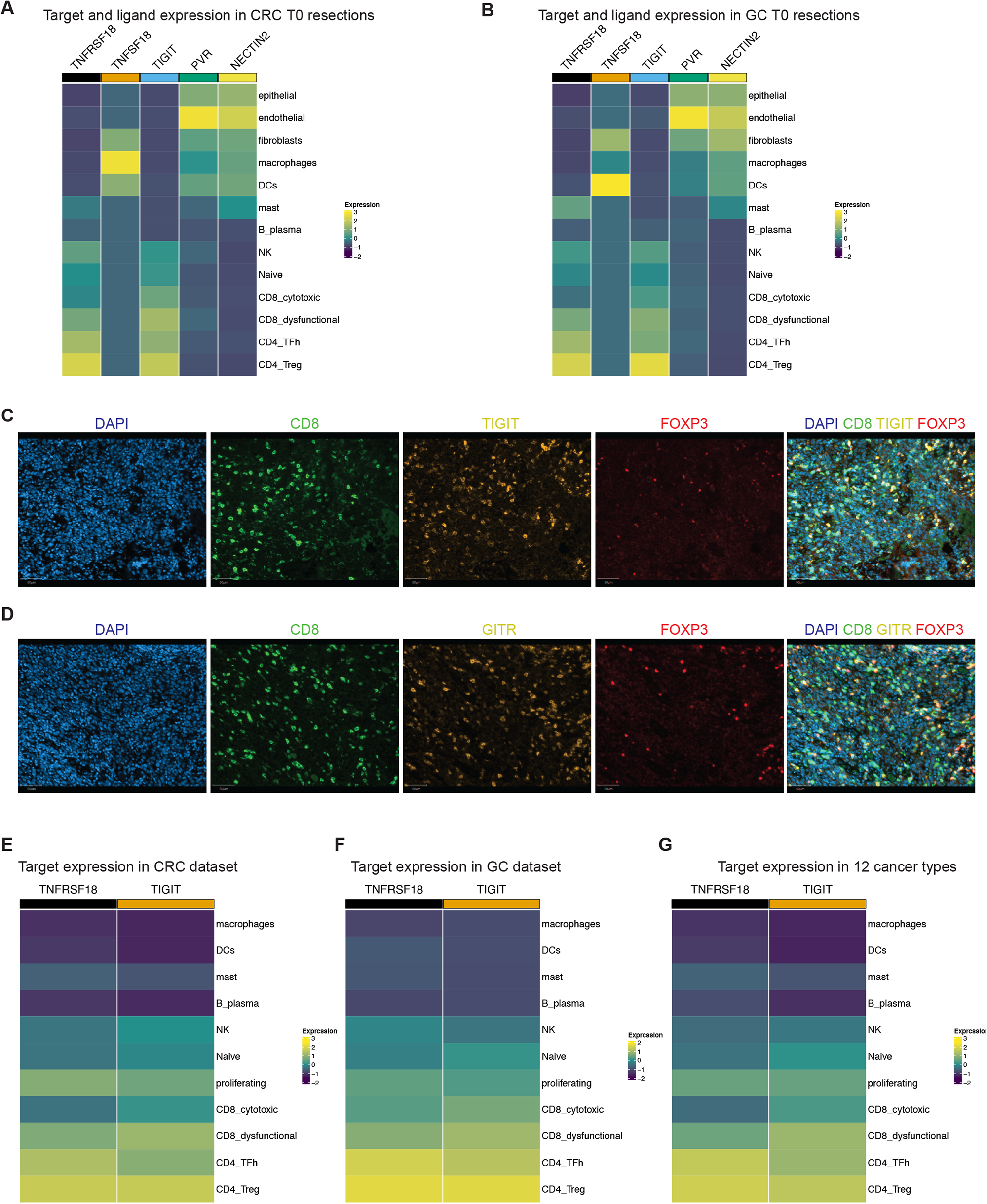
(A-B) Scaled expression of respective genes in various cell types from (A) all CRC T0 resections, (B) all GC T0 resections. (C-D) Immunofluorescence staining for respective proteins or their merged image in an example region of interest from sample CRC-2. Scale bar = 50 μm. (E-G) Scaled expression of respective genes in various cell types from (E) CRCs in the publicly available tumor immune atlas dataset, (F) our previously published GC dataset and (G) remaining 12 tumor types in the tumor immune atlas dataset.

We also measured the expression of GITR and TIGIT proteins in these baseline tumor tissues (T0 resections) using multiplexed immunofluorescence staining. We stained the tumors using two independent antibody panels containing CD8, FOXP3 and TIGIT or CD8, FOXP3 and GITR respectively (**Fig. 2C, D**). We performed image analysis using a multiplex classifier for detecting single stain or double stain positive cells as described previously (17). From all samples, an average of 37.4% of total CD8 positive cells expressed TIGIT. An average of 53.68% of total FOXP3 positive cells were TIGIT positive (**Supplemental Fig. 1B, C**). Similarly, 42.46% of CD8 cells expressed GITR (**Supplemental Fig. 1D, E**). Among the FOXP3 cells, 74.5% expressed GITR. These results confirmed that among our tumor samples, CD8 T cells and Tregs expressed the TIGIT and GITR protein.

Overall, these results indicated that expression of our target receptors and their ligands was prevalent in the TME of the harvested cancers. Targeting these receptors has the potential to modify the function of anti-tumor T cell subsets such as cytotoxic CD8, dysfunctional CD8 and TFh-like cells, as well as immunosuppressive Tregs.

### Expression of TIGIT and GITR in colorectal and other cancer types

To determine the expression of these two targets among an independent and expanded set of colorectal, gastric and other tumor types, we analyzed gene expression among a data set of 13 different cancer types (21). Importantly, this dataset included 25 independent CRCs. Other cancer types included breast carcinomas (**BC**), basal cell and squamous cell carcinomas (**BCC**), endometrial adeno-(**EA**) and renal cell carcinomas (**RCC**), intrahepatic cholangiocarcinoma (**ICC**), hepatocellular carcinomas (**HCC**), pancreatic ductal adenocarcinomas (**PDAC**), ovarian cancers (**OC**), non-small-cell lung cancers (**NSCLC**), and cutaneous (**CM**) and uveal (**UM**) melanomas. For gastric cancer, we evaluated our previously published dataset of seven GC samples (17). High *TNFRSF18* and *TIGIT* expression was detected in dysfunctional and cytotoxic CD8 T, TFh-like, Treg and proliferating cells (**Fig. 2E-G**). Overall, this result confirmed the expression of these targets in additional CRC and GC tumors. These targets are also expressed in a wide variety of solid tumor types.

### Primary tissue slice cultures maintain the native TME composition of gastrointestinal cancers

As noted previously, tissue slice cultures have been demonstrated to maintain a high degree of tissue viability, cellular diversity, and cellular transcriptional profiles (6, 10). We confirmed that the TSCs cultured for 24 hours maintained the cellular characteristics similar to the baseline state of the TME (i.e., T0 cell conditions at time of resection). As noted previously, other groups have used short culture periods to maintain the cellular diversity found in the native TME (4–7). First, we evaluated the TSC cellularity using hematoxylin and eosin (**H&E**) staining of the cultures. This result showed that cell morphology remained intact with little evidence of necrosis or other signs of overt cell death (**Supplemental Fig. 1F)**.

Next, we evaluated the single cell gene expression across the two conditions for all samples which included: i) the baseline T0; ii) TSC post-24 hour incubation with isotype control antibody; (**Fig. 3A**). We corrected the data for experimental batch but not for the experimental condition, using the Harmony algorithm (24). Cells belonging to the baseline T0 tissue and control clustered together in the Uniform manifold approximation and projection (**UMAP**). This result indicated that the cells had similar gene expression profiles. We calculated the Adjusted Rand Index (**ARI**) to determine the variation in gene expression between the baseline and control culture. Cluster labels compared to the experimental condition had a low ARI value of 0.009 which indicated that clustering was due to the cells having similar gene expression characteristics and not driven by the T0 or TSC experimental condition.

**Figure 3.**
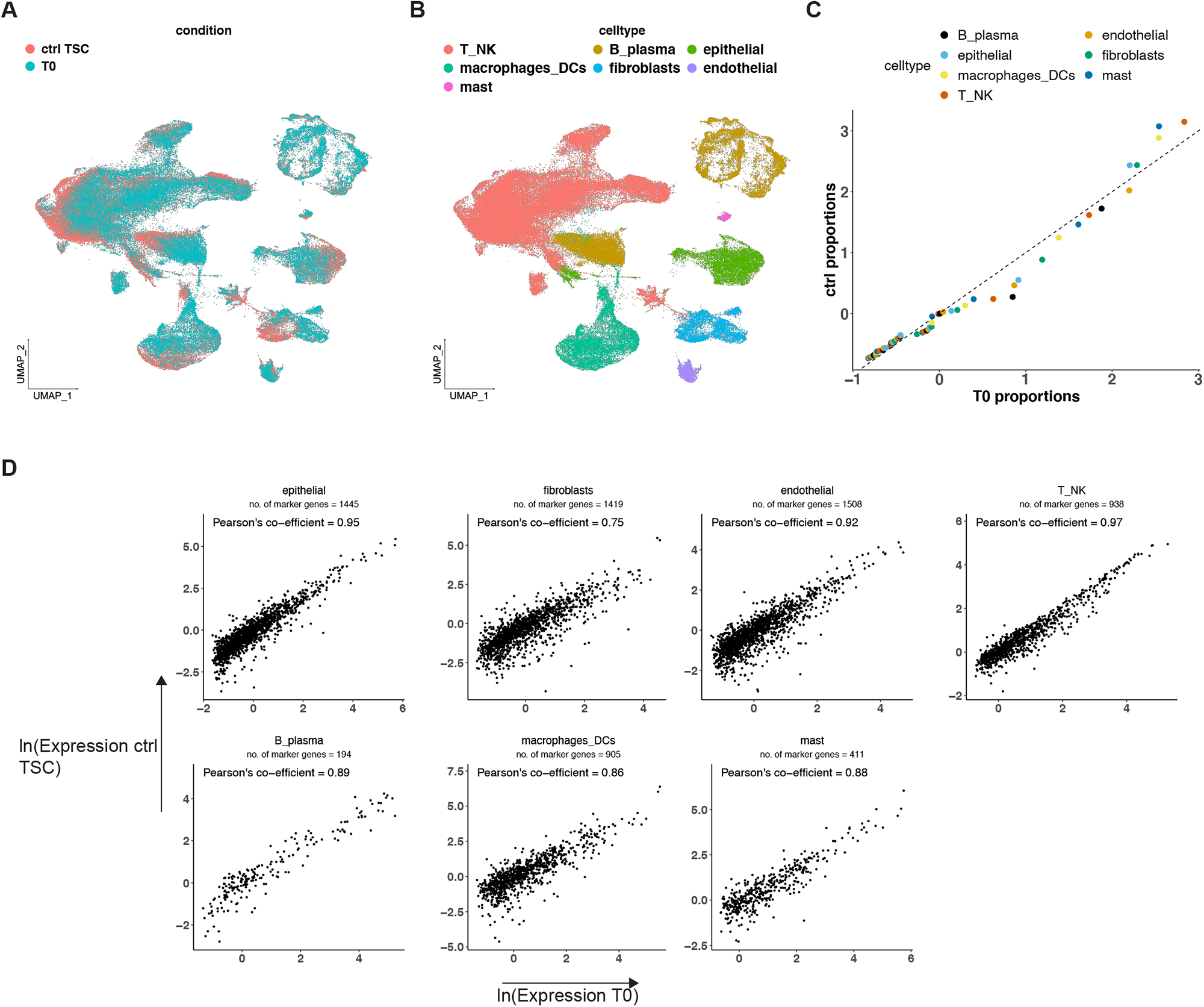
(A-B) UMAP representation of dimensionally reduced data from T0 and 24 hour ctrl TSCs following batch-corrected graph-based clustering of all datasets colored by (A) experimental condition and (B) cell type. (C) Quantile-quantile plot comparing the proportion distributions of respective cell lineages across all T0 and ctrl TSCs. (D) Scatter plot indicating average log expression of marker genes for T0 cell lineages in T0 and ctrl TSC in respective cell lineage, annotated with the number of marker genes examined. Pearson’s co-efficient was calculated using non-log transformed values.

We annotated cell types for each cluster using marker gene expression as previously described. The TSC samples contained all cell lineages compared to the matched baseline T0 samples. These cell types included tumor epithelial, macrophages and dendritic cells, T, NK, B or plasma lymphocytes, mast cells, fibroblasts, and endothelial cells (**Fig. 3B**). The relative proportion of all cell lineages was also maintained in the TSCs compared to the baseline tumor tissue (**Fig. 3C**).

We identified marker genes for each cell lineage in the baseline T0 samples (Seurat Wilcoxon test, log fold change >= 0.25, adjusted *p* <= 0.05). We compared the average gene expression of these marker genes in each respective cell lineage in T0 to TSCs. Expression was highly correlated across all cell types (**Fig. 3D**) (Pearson correlation >=0.75, *p* <= 2.63E-67). Hence, the TSCs maintained the cellular heterogeneity and cell states that were present in the original tumor.

### General stimulation of T cells and other cell types in the TSC TME

To demonstrate that the TSC cells were functionally responsive, we used a stimulus with phorbol ester 12-myrisate 13-acetate and the calcium ionophore ionomycin (**PMA/Ionomycin**). In combination, these compounds stimulate downstream pathways associated with T cell activation (31). We integrated data from all experiments and performed cell type identification using marker based and SingleR assignments (**Methods**). We evaluated specific subsets of cells from samples treated with ctrl and each respective perturbation. To account for inter-patient variability in differential expression (**DE**) analysis, we utilized model-based analysis of single-cell transcriptomics (**MAST**) (32) incorporating sample as a random effect in the model (33). A threshold of log fold change of 0.25 and false discovery rate (**FDR**) *p* < 0.05 was used to identify significantly DE genes.

From the TSCs exposed to PMA/Ionomycin, we detected differentially expressed genes associated with activation in CD8 T cells (*CD69, CRTAM*) (**Fig. 4A**), TFh-like cells (**Fig. 4B**) (*CD69, CD40LG, IL2RA*) and Tregs (*CTLA4, TNFRSF4, TNFRSF9, IL2RA*) (**Fig. 4C**). All cell types also responded with increased expression of NR4A and EGR family genes which are associated with signaling of the nuclear factor of activated T cells (**NFAT**). This pathway is associated with T cell activation and anergy following stimulation (34). In CD8 T and TFh-like cells, we also identified increased expression of several effector cytokines and chemokines including *CCL4*, *CCL3*, *IFNG*, *TNF*, etc. relative to control. At the pathway level, we confirmed a significant increase in NF-KB signaling and calcium ion response pathway activity in CD8 T cells (**Fig. 4D, E**). Both pathways are known mediators of effects of PMA/Ionomycin (31). Across all the tumors, CD8 T cells consistently responded to PMA/Ionomycin stimulation as indicated by a significant increase in NF-KB activity (**Fig.4F**).

**Figure 4.**
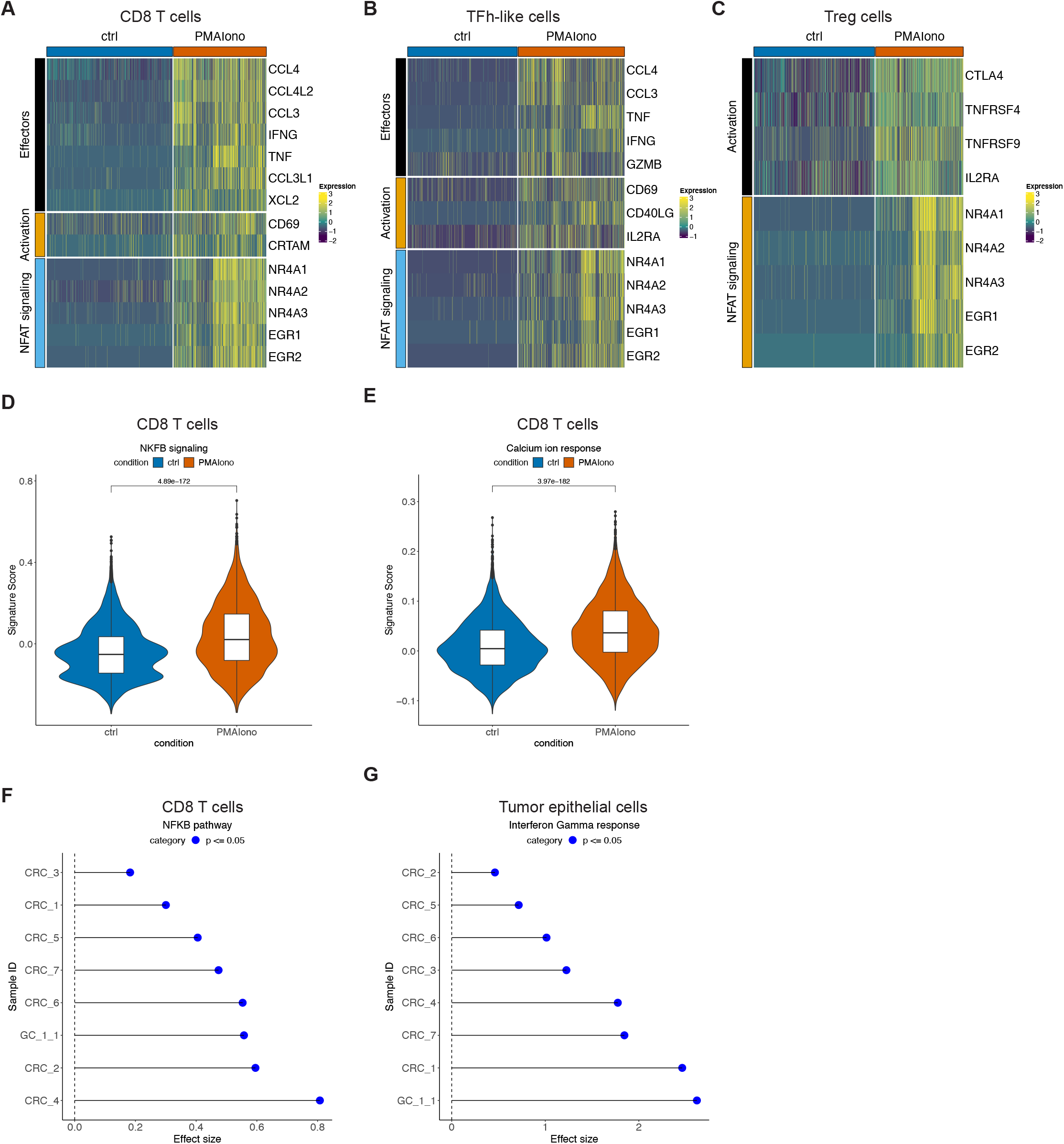
(A-C) Scaled expression of respective genes in control or PMA/Ionomycin treated (A) CD8 T cells, (B) TFh-like cells and (C) Treg cells. (D-E) Respective pathway activity in control and treated CD8 T cells with T-test *p*. (F-G) Cohen’s effect size and *p* of t-test comparison of respective pathway activity between control and treated cells from each individual sample.

The tumor epithelial cells showed significant increases in interferon (**IFN**) gamma response – signaling across all tumor samples (**Fig. 4G**). This indicated that increased IFN from activated T cells was able to affect neighboring tumor cells, reflecting the preserved intercellular networking in TSCs. Overall, these experiments with PMA/Ionomycin confirmed that the TSC cells were functional and demonstrated intercellular interworking among the cells in the TSCs.

### GITR activation had limited and heterogenous effects on CD8 T cell cytotoxicity

We evaluated the effects of the GITR agonist on the TSCs. For CD8 T cells, the only gene which showed differential expression across all nine tumors was *CCL4*. This gene had a fold change of >0.25 upon GITR agonist treatment (MAST DE) (**Fig. 5A**). Other significantly increased genes with lower fold changes (>0.15, FDR < 0.05) included cytokines *CCL4L2*, *GZMA*, *GNLY*, *CCL3* and *PRF1* (**Supplemental Table 2**). Thus, GITR agonist exposure had limited effects on gene expression across the tumors.

**Figure 5.**
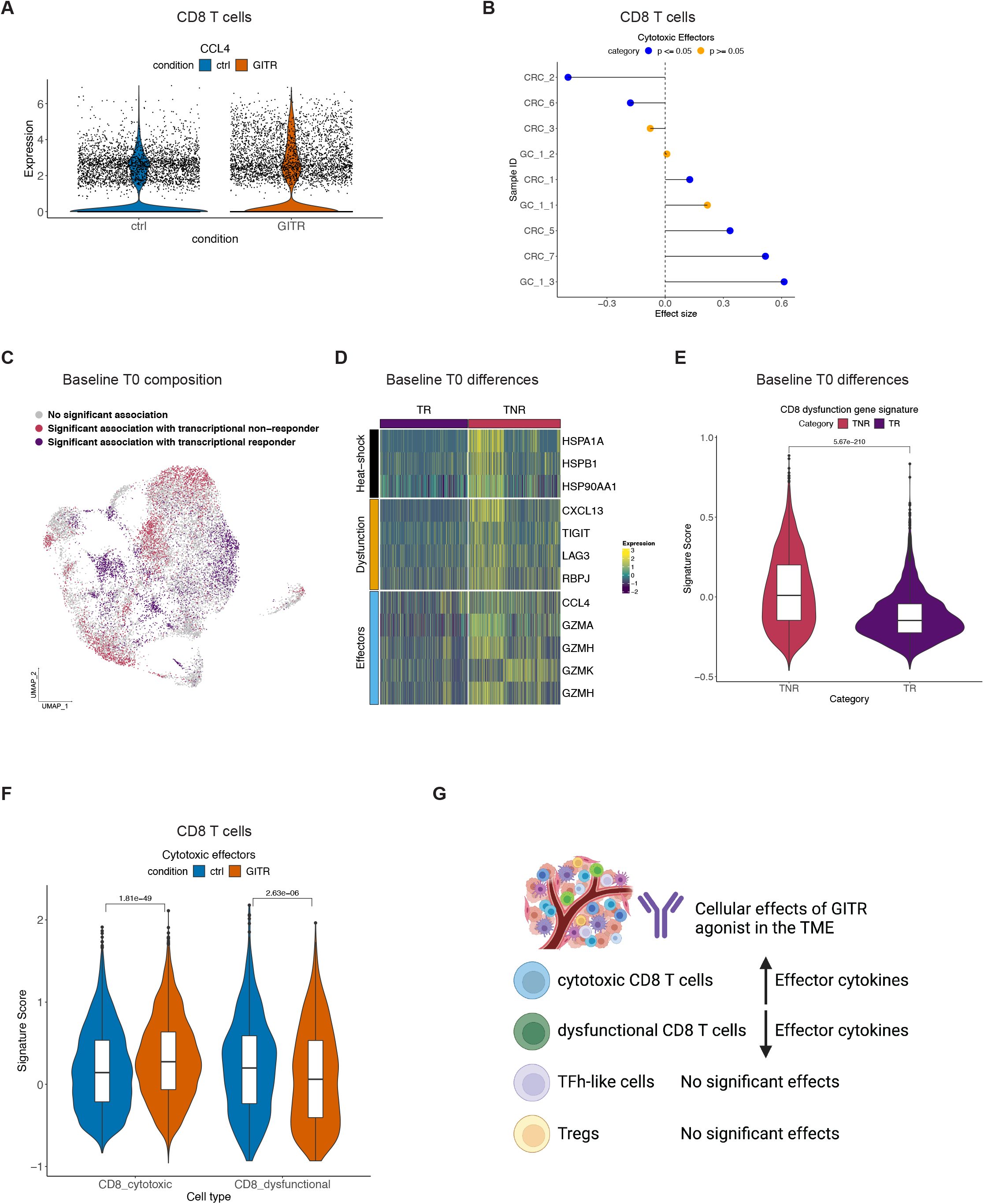
(A) Violin plot depicting the expression of *CCL4* in control or GITR agonist treated CD8 T cells derived from all samples. (B) Cohen’s effect size and *p* of t-test comparison of cytotoxic effector pathway activity between control and treated CD8 T cells from each individual sample. (C) UMAP representation of T0 samples identifying cells significantly associated with transcriptional responders (TR) or non-responders (TNR) based on differential abundance analysis. (D) Scaled expression of respective genes in cells significantly associated with TR or TNR. (E) Expression of gene signature of CD8 T cell dysfunction in TR and TNR with t-test *p*. (F) Cytotoxic effector pathway activity in control and treated cytotoxic and dysfunctional CD8 T cells with t-test *p*. (G) Schematic representation summarizing the *ex vivo* effects of GITR agonist in the TME.

We evaluated the effect of GITR agonist on the expression of a CD8 T cell cytotoxic gene expression signature that has been previously reported (35, 36) (**Supplemental Table 3**, **Fig. 5B**). Significant increases in cytotoxic gene signature expression were observed in four (CRC-1, GC-1-3, CRC-5, CRC-7) out of nine tumors. Hence, the GITR agonist showed interpatient variability in terms of a gene expression response.

### Dysfunctional CD8 T cells were indicators of no response to the GITR agonist

We investigated the differences in transcriptional responsiveness to GITR agonist. GITR agonist exposure led to only a limited increase in cytotoxic effectors for only a subset of the tumors. We grouped tumors as transcriptional responsive (**TR**) when they responded with an increase in cytotoxic effector gene expression upon treatment (GC-1-3, CRC-1, CRC-5, CRC-7). Tumors lacking this response were identified as transcriptional non-responsive (**TNR**) (GC1-1, GC-1-2, CRC-3, CRC-2, CRC-6). We compared the baseline CD8 T cells (T0) in the TNR versus TR samples. We used differential abundance (**DA**) analysis (37) to identify cell populations with significantly different distributions based on the CD8 T cells’ gene expression. We identified cells in the UMAP representation that were significantly associated with either TNR or TR status (**Fig. 5C**) (Wilcoxon *p* <= 0.065) and identified differentially expressed genes between the two.

The TNR associated cells had significantly increased expression (Seurat Wilcoxon adjusted *p* < 0.05, log fold change >= 0.25) of CD8 T cell dysfunction markers including *CXCL13*, *TIGIT*, *LAG3* and *RBPJ* (19) (**Fig. 5D**). Additionally, these cells had increased expression of effectors including the granzyme family genes. These cells also had increased expression of HSP family genes that have been linked to exhaustion (38). Also, we evaluated a gene signature for CD8 T cell dysfunction that has previously been described (26) (**Supplemental Table 3**). TNR associated cells had significantly higher levels of dysfunction (**Fig. 5E**). This result indicated that CD8 T cells with this dysfunctional phenotype do not respond to GITR agonist.

To test this association between a lack of transcriptional response in dysfunctional CD8 T cells, we examined *ex vivo* responses among CD8 T cell subsets across all tumors (**Fig. 5F**). Increased effector gene signature upon treatment was restricted to cytotoxic CD8 T cells. In dysfunctional cells, GITR agonist reduced effector gene expression. Hence, GITR agonist only stimulated effector cytotoxic cells. However, in exhausted dysfunctional cells this stimulation reduced the cytotoxic potential.

Among TFh-like cells, the GITR agonist led to a significant increase in gene expression of only three genes (*S100A4*, *MT-CO1*, *MT-ND1*). No significant changes were detected in Tregs and NK cells. These results indicated a limited effect of GITR agonist in the TME (**Fig.5G**).

### TIGIT inhibition activated CD8 T cells in the TME

Next, we evaluated the effects of the TIGIT antagonist on the TSCs of five tumors (CRC-4, CRC-5, CRC-7, GC-1-2, GC-1-3). After TIGIT antagonist exposure, CD8 T cells showed increased expression of several cytotoxic effector genes (**Supplemental Table 4**, **Fig.6A**). These genes included *IL32*, the granzyme family genes, *PRF1*, *NKG7*, *CCL5* and *CCL4*. We identified increased expression of genes involved in actin cytoskeleton remodeling including *PFN1*, *COTL1* and *CORO1A*. The *CD3D* and *CD3DE* genes increased upon TIGIT inhibition. These genes are components of TCRs. Overall, TIGIT inhibition increased TCR signaling and activation of CD8 T cells.

**Figure 6.**
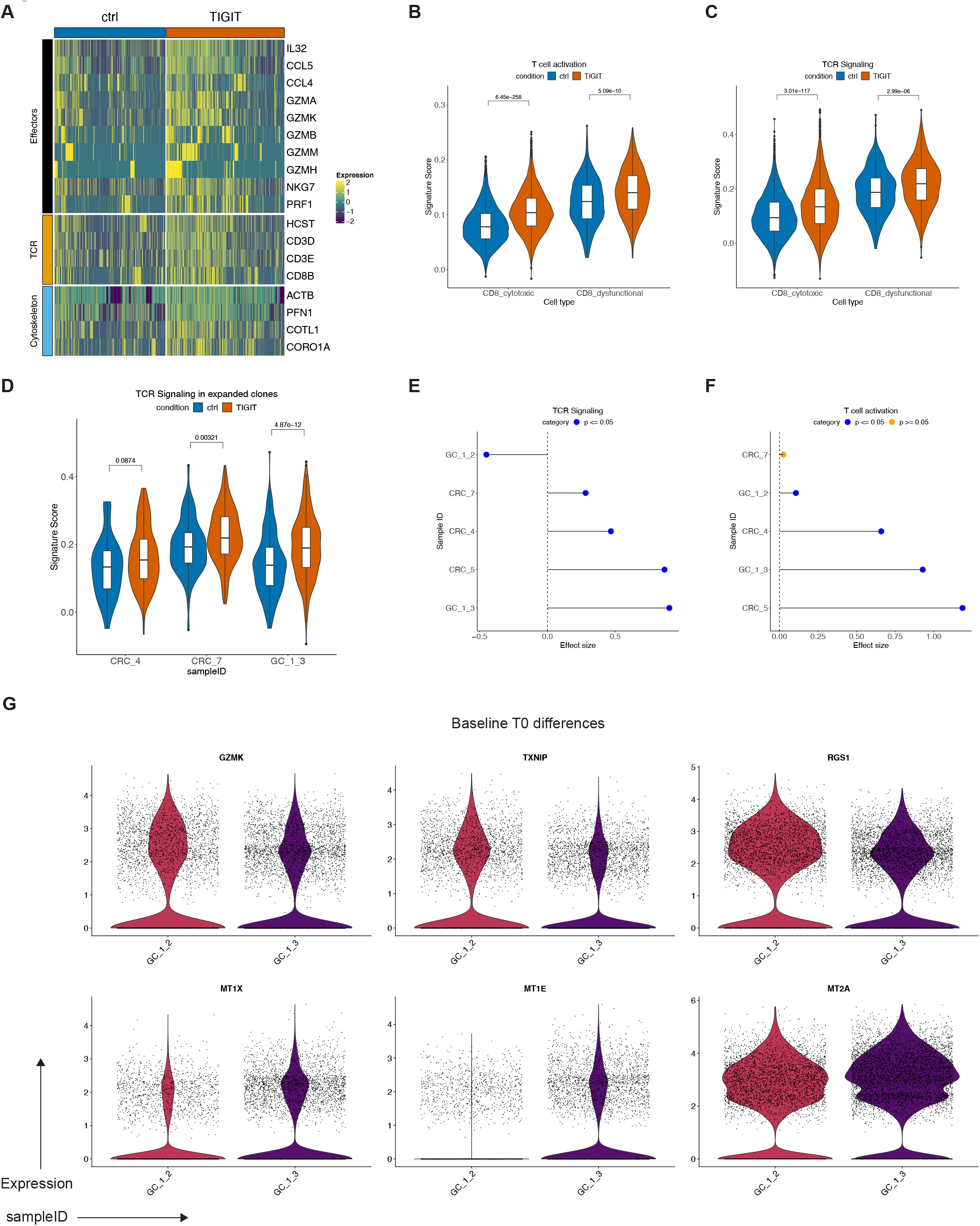
(A) Scaled expression of respective genes in control or TIGIT inhibitor treated CD8 T cells. (B-D) Respective pathway activity in control and treated CD8 T cells with t-test *p* in (B-C) CD8 T cell subtypes and (D) baseline expanded CD8 TCR clonotypes per sample. (E-F) Cohen’s effect size and *p* of t-test comparison of respective pathway activity between control and treated cells from each individual sample. (G) Violin plots depicting the expression of respective genes in CD8 T cells from GC-1-2 and GC-1-3 samples.

We evaluated the effect of TIGIT antagonist among the different CD8 cell subtypes (i.e., cell states) by quantifying TCR signaling and T cell activation pathways. We observed significantly increased TCR signaling and T cell activation in both cytotoxic and dysfunctional CD8 T cells (**Fig. 6B, C**). This result indicated that TIGIT inhibition is capable of reinvigorating dysfunctional exhausted cells. In contrast, the GITR agonist reduced the cytotoxicity of dysfunctional cells.

### TIGIT inhibition activated specific CD8 clonotypes

TIGIT inhibition had specific effects on certain CD8 TCR clonotypes. From the baseline tumor tissue (T0), we identified TCR clonotypes in CD8 T cells across patients as previously described. Clonotypes that were present in more than one cell were indicative of potential tumor reactivity (**Fig. 1E**) (2). We used these TCR clonotypes from the baseline to identify how the CD8 T cells with the same clonotype responded to TIGIT antibody versus the control in the TSCs. In three tumors, we recovered 14-28% of these clonotypes in both the ctrl and TIGIT conditions, allowing us to examine the effect of treatment in these cells. TIGIT inhibition successfully increased TCR signaling among these clones (**Fig. 6D**). This result indicated that TIGIT treatment can specifically increase the activation of potential anti-tumor clonotypes.

### TIGIT inhibition generates a variable cellular response in a metastatic gastric cancer

Using these gene signatures of TCR signaling or T cell activation, we examined tumor-specific responses to TIGIT inhibition. The CD8 T cells from all tumors responded with a significant increase in either one or both processes of TCR signaling and T cell activation upon treatment (**Fig. 6** **E, F**). We validated the significant increase in expression of downstream cytotoxic effector *GZMB* using RNA in situ hybridization (**RNA-ISH**) in tumors CRC-5 and GC-1-3 (**Supplemental Fig. 2A, B**). Hence, TIGIT inhibition activated local infiltrating CD8 T cells in the TME across all tumors.

A notable response pattern was observed for tumors GC-1-2 and GC-1-3 (**Fig. 6** **E, F**). The GC-1 tumors samples represented patient matched pairs of peritoneal metastases (**Table 1**). Interestingly, there was a variation in the response among these two metastases. Compared to GC-1-3, tumor GC-1-2 had a lower increase in the extent of T cell activation and responded with a decrease in TCR signaling upon treatment (**Fig. 6E, F**). This indicated a reduced responsiveness to TIGIT inhibition in CD8 T cells in GC-1-2 compared to GC-1-3.

To determine the factors leading to variation in transcriptional response from two metastatic tumors, we examined differences in their baseline T0 CD8 phenotypes. GC-1-2 CD8 T cells had significantly higher expression of *GZMK* (**Fig. 6G**) associated with effector memory CD8 cells (2) and *RGS1* associated with pre-exhausted and exhausted CD8 T cells (39). The reduced responsive cells also had upregulated *TXNIP*, which has been demonstrated to reduce effector functions in CD8 T cells in viral infection (40). Conversely, GC-1-3 had increased expression of metallothionein genes *MT1E*, *MT1X* and *MT2A*. In a recent study, metallothionein family genes were demonstrated to link levels of CD8 activation and dysfunction to modulate their effector capacity (41). In summary, we identified that TIGIT inhibition had different effects across two metastatic gastric cancers from the same patient. Variation in TIGIT response was associated with genes that modulate effector, activation, and dysfunctional phenotypes among CD8 T cells.

### TIGIT inhibition activated TFh-like cells in the TME

TIGIT inhibition led to activation of TFh-like cells, a cellular effect that has not been described previously. Differential expression analysis identified the upregulation of *ACTB*, *PFN1*, *S100A4*, *S100A6* and *TAGLN2* that are involved in T cell activation (42) (**Supplemental Table 5,** **Fig. 7A**). These cells also upregulated *IL32* expression, a cytokine with potential proinflammatory effects. Expression of *IL32* in the TME has been associated with response to PD-1 inhibition (43). Importantly, TIGIT inhibition led to the increased expression of *CXCL13*. TFh-like cells which express CXCL13 may be associated with B cell response and generation of tertiary lymphoid structures (44). These features mediate an effective immune response against a tumor. These effects were confirmed at the pathway level where TIGIT antagonist treatment led to a significant increase in the T cell activation ontology program (**Fig. 7B**). This effect was observed in four (CRC-4, CRC-5, GC-1-2, GC-1-3) out of five patients (**Fig. 7C**).

**Figure 7.**
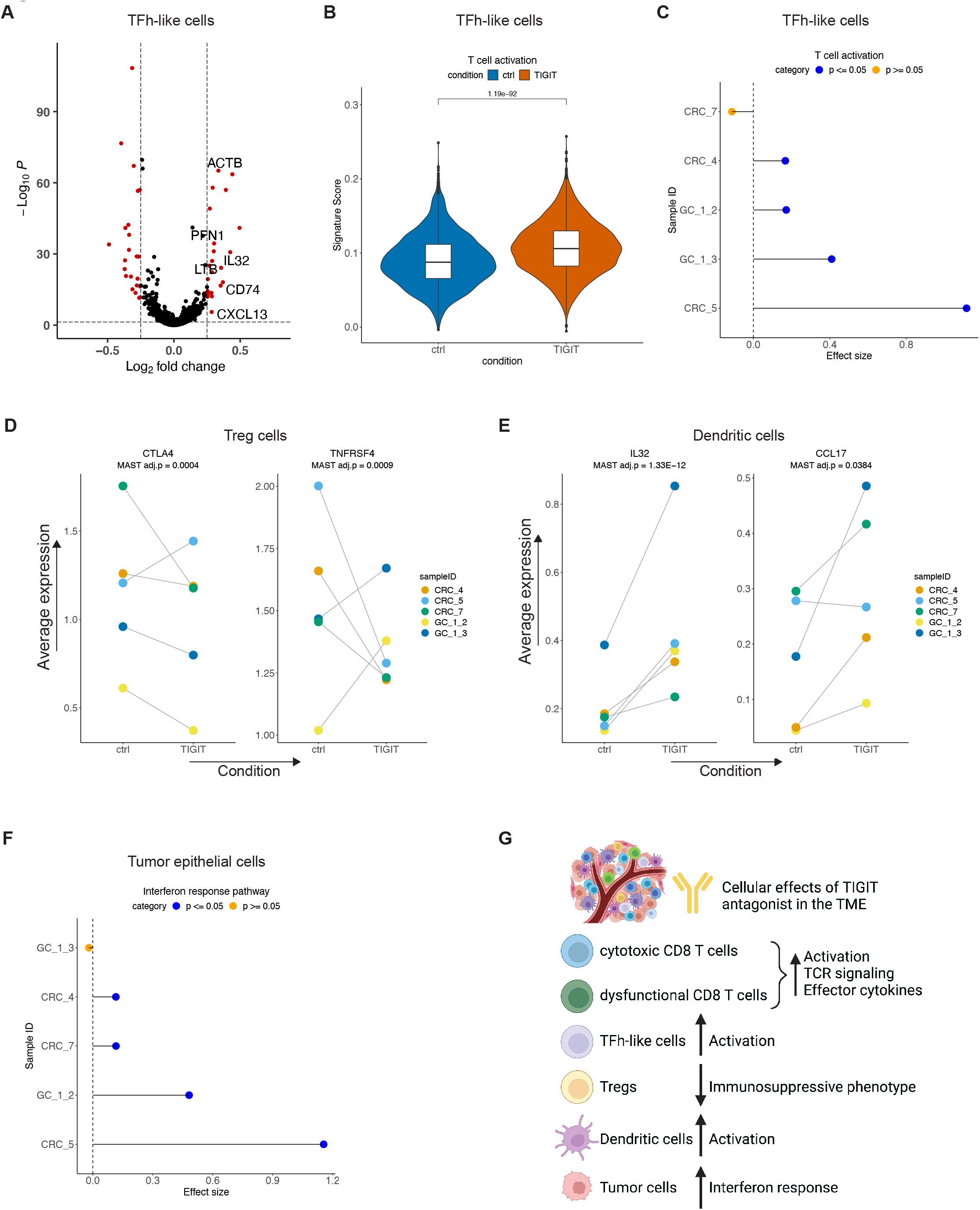
(A) Volcano plot demonstrating significant differentially expressed genes in TFh-like cells relative to control following TIGIT inhibition. (B) Pathway activity in control and treated TFh-like cells with t-test *p*. (C) Cohen’s effect size and *p* of t-test comparison of pathway activity between control and treated TFh-like cells from each individual sample. (D) Average expression of respective genes in each sample in control and treated Treg cells with MAST DE adjusted p-value. (E) Average expression of respective genes in each sample in control and treated dendritic cells with MAST DE adjusted p-value. (F) Cohen’s effect size and *p* of t-test comparison of pathway activity between control and treated tumor epithelial cells from each individual sample. (G) Schematic representation summarizing the *ex vivo* effects of TIGIT antagonist in the TME.

Increased TFh-like cells have been demonstrated to predict response to and are proposed to be a target of PD-1 immunotherapy (45, 46). However, the effects of targeting these cells in the human TME have remained unknown. We demonstrated that TIGIT antagonist activated these cells in the local TME. This represented an important cellular mediator of response to TIGIT inhibition that can generate an inflammatory anti-tumor TME.

### TIGIT inhibition’s effects on other cell types in the TME

The TIGIT antibody had notable effects among the Treg cells (**Supplemental Table 6**). We observed an increase in *CST7*, which is associated with TCR signaling (47), and *CD7* linked to a mature Treg phenotype (48). However, this was accompanied by a reduction in *CTLA4* and *TNFRSF4* expression. A reduction in expression in either of these two genes was seen in all patients (**Fig. 7D**). Both these molecules are key regulators of an immunosuppressive Treg phenotype (49). This indicated modest effects of TIGIT inhibition on Tregs with a reduction in immunosuppressive phenotype.

TIGIT inhibition impact on DCs included a significant increase in expression of *CCL17* and *MARCKSL1* – these genes are indicators of DC activation and a maturation phenotype (50). Accompanying this activated DC phenotype, was a significant increase in *IL32* expression (**Supplemental Table 7,** **Fig. 7E**). DCs which express IL32 can activate T cell responses in the TME (51). Conversely *IL2RA*, which is associated with an immunosuppressive DC phenotype (52), was reduced with TIGIT inhibition. TIGIT antagonist thus led to activation of DCs in the TME. This result has potential implications in improving antigen presentation and T cell priming to orchestrate an anti-tumor response in the TME.

Exposure to the TIGIT antagonist did not lead to any gene expression changes among NK cells. Finally, we examined the effects in tumor epithelial cells. TIGIT treatment led to an increase in IFN response signature among four (CRC-4, CRC-5, CRC-7, GC-1-2) out of five tumors (**Fig. 7F**). Overall, these results indicated that the proinflammatory effects modulated by TIGIT in various cell types in the TME translated into initial favorable transcriptional responses at 24 hours in tumor epithelial cells. This included activation of both cytotoxic and dysfunctional CD8 T, TFh-like cells and DCs together with a reduced immunosuppressive phenotype in Tregs, which can promote a favorable inflammatory TME (**Fig. 7G**).

## DISCUSSION

Many immunotherapy agents and combinations are being studied in clinical trials, often with disappointing results (53). It is important to determine the cellular basis for how these agents work in a dynamic and complex TME. An analysis of the TME and its response to these agents is critical to prioritize targets, identify mechanisms of resistance and design rational treatment combinations. Our experimental design combined a robust culture system that preserves the original TME together with single-cell readouts that provide granular insights into the mechanism of action of perturbations. This identified heterogenous cellular and patient responses to GITR and TIGIT immunotherapy in the TME of GC and CRC.

Despite promising results from T cell culture, mouse, and primate models, GITR agonists have shown no meaningful clinical responses in recent clinical trials (54–58). Our results demonstrated that GITR agonist has limited activity in the TME, restricted to cytotoxic CD8 T cells that lack exhaustion features. Moreover, dysfunctional cells had a decrease in cytotoxic activity upon GITR stimulation. Given that PD-1 inhibitors act to re-invigorate exhausted CD8 T cells, our finding raises the possibility that combining them with GITR agonists might antagonize this effect. We also saw no effects on Treg reprogramming in the TME with GITR agonist. In clinical trials, GITR agonist mediated depletion of Tregs in the peripheral blood or TME has been observed only in some patients (58).

Compared to GITR agonist, we saw widespread effects in different cell types in the TME with TIGIT inhibition. This included activation of CD8 T cells and TFh-like cells. Both these components can mediate anti-tumor immunity. Our observation that TIGIT inhibition can increase TCR signaling in expanded CD8 clonotypes suggests that tumor antigen specific T cells could potentially be reinvigorated with treatment. Early reports have demonstrated some clinical responses with TIGIT monotherapy or in combination with PD-1 (59). However, these responses are likely to be restricted to only a subset of patients (60). We did observe variation in the extent of transcriptional responses in CD8 T cells in our samples, which was associated with differential baseline expression of *GZMK*, *RSG1*, *TXNIP* and metallothionein family genes. An expanded study with greater number of samples and mechanistic studies will allow us to examine these correlates of response to TIGIT inhibition.

TSCs remain viable for 1-2 weeks in culture (5). We evaluated the short-term perturbation effects after exposure to specific antibodies. This feature enables culture in media free from cytokines such as IL-2 that are routinely used in maintaining T cells in culture for longer duration. As we have demonstrated previously, IL-2 can reprogram transcriptional T cell states (61). Our approach allows the evaluation of cell states in the original TME. Most studies of immunotherapy agents lack this feature.

A limitation of TSCs is that they allow interrogation only of the local TME, but not of peripheral and lymph node immune responses. These elements are also important players in the clinical response to immunotherapy (62). Short term readouts also do not capture remodeling of the TME that could occur over longer duration. However, a recent study demonstrated that short term fragment culture responses to PD-1 were corelated with long term clinical responses (3). While it only assesses the local TME response, our experimental strategy fills an important gap in preclinical studies.

Overall, our study identified cellular mechanisms of action of GITR and TIGIT immunotherapy in the TME of patients. Future studies testing combination therapies with PD-1, targeting macrophages and fibroblasts in the TME and examining the clinical predictive value of *ex vivo* responses will further improve clinical translation.

## ONLINE METHODS

### Sample acquisition

This study was conducted in compliance with the Helsinki Declaration. All patients were enrolled according to a study protocol approved by the Stanford University School of Medicine Institutional Review Board (IRB-44036). Written informed consent was obtained from all patients. All samples were surgical tumor resections. Clinical pathology report that was generated at the time of the resection was reviewed for all samples.

### Tissue processing

Tissues were collected in plain RPMI on ice immediately after resection and dissected with iris scissors. From the original T0 surgical resections, a portion was fixed for histopathology, a portion was subjected to dissociation and the remainder was used to generate tumor slice cultures.

### *Ex vivo* tumor slice cultures (TSCs)

A VF-310-0Z Compresstome tissue slicer and its accessories (Precisionary, Greenville, NC, USA) were used to generate tissue slices from a piece of the resection. Tissue sample was glued onto the specimen tube base using All Purpose Krazy Glue (Elmer’s Products, Inc., Westerville, OH, USA). The 3% agarose solution was prepared by diluting UltraPure Low Melting Point Agarose (ThermoFisher Scientific) in water followed by heating in a microwave and cooling for around three minutes at room temperature. The tissue sample was retracted into the specimen tube and covered with agarose solution. Agarose was solidified by placing pre-chilled chilling block supplied by manufacturer over the specimen tube. Specimen tube was assembled onto compresstome as per manufacturer’s instructions and cold PBS was used as a solution in the buffer tank. Slices were generated using advance setting of 3, oscillation of 5 and thickness of 400 μm. Slices were placed onto a 0.4 μm pore size Millicell Cell Culture Insert (Sigma-Aldrich, St. Louis, MO, USA) that was then placed into a 35-mm dish (ThermoFisher Scientific). The media volume included 1.5 ml that was placed into the surrounding dish and 0.5 ml placed onto the slices followed by culture in a cell culture incubator. Media was composed of RPMI, 10% FBS, 1% Antibiotic-Antimycotic (ThermoFisher Scientific). Perturbations were added to the media once at the beginning of culture. 2 μg/ml IgG1 Fc (BPS Bioscience, catalog #71456) was used as control. Treatment conditions included 2 μg/ml GITR agonist (BPS Bioscience, catalog #79053), 2 μg/ml TIGIT antagonist (BPS Bioscience, catalog #71340) or 6 μg/ml eBioscience Cell Stimulation Cocktail (500X) (ThermoFisher Scientific). At 24 hours, TSCs were subjected to fixation for histology and dissociation.

### Histopathology

Tissue was fixed in 10% formalin for approximately 24 hours at room temperature. Paraffin embedding and hematoxylin and eosin staining was conducted by the Human Pathology Histology Services core facility at Stanford University. Whole slide images were obtained using Aperio AT2 whole slide scanner (Leica Biosystems Inc., IL, USA). Tissue fixation was not performed for T0 sample CRC-3 due to inadequate material.

### Single-cell dissociation

Tissue dissociation was conducted using a combination of enzymatic and mechanical dissociation using a gentleMACS Octo Dissociator (Miltenyi Biotec) as described previously (63). Cells were cryofrozen using 10% DMSO in 90% FBS (ThermoFisher Scientific, Waltham, MA) in a CoolCell freezing container (Larkspur, CA) at -80 °C for 24-72 hours followed by storage in liquid nitrogen. For scRNA-seq, cryofrozen cells were rapidly thawed in a bead bath at 37 °C, washed twice in RPMI + 10% FBS, and filtered successively through 70 μm and 40 μm filters (Flowmi, Bel-Art SP Scienceware, Wayne, NJ). Live cell counts were obtained using 1:1 trypan blue dilution. Cells were concentrated between 500-1500 live cells/μl.

### Single-cell RNA sequencing

The scRNA-seq libraries were generated from cell suspensions using Chromium Next GEM Single Cell 5’ version 1.1 (samples CRC-1, CRC-2, GC1-1, GC-1-2, GC-1-3) or version 2 (samples CRC-3, CRC-4, CRC-5, CRC-6, CRC-7) (10X Genomics, Pleasanton, CA, USA) as per manufacturer’s protocol. All libraries from a patient were prepared in the same experimental batch. Ten thousand cells were targeted with 14 PCR cycles for cDNA and library amplification. Chromium Single Cell V(D)J Human T Cell Enrichment Kit was used to prepare TCR libraries from single-cell cDNA as per manufacturer’s protocol. A 1% or 2% E-Gel (ThermoFisher Scientific, Waltham, MA, USA) was used for quality control evaluation of intermediate products and sequencing libraries. Qubit (Thermofisher Scientific) was used to quantify the libraries as per the manufacturer’s protocol. Libraries were sequenced on Illumina sequencers (Illumina, San Diego, CA).

### Data processing of scRNA-seq

Cell Ranger (10x Genomics) version 3.1.0 or 5.0.0 ‘mkfastq’ command was used for NextGEM version 1.1 and version 2 libraries respectively to generate Fastq files. Cell Ranger version 3.1.0 ‘count’ was used with default parameters and alignment to GRCh38 to generate a matrix of unique molecular identifier **(UMI)** counts per gene and associated cell barcode. Cell Ranger version 6.0.0 ‘vdj’ command was used to perform sequence assembly and clonotype calling of TCR libraries with alignment to the prebuilt Cell Ranger V(D)J reference version 5.0.0 for GRCh38.

### Clustering individual datasets

We constructed Seurat objects from each sample using Seurat (version 4.0.1) (64, 65). We applied quality control filters to remove cells that expressed fewer than 200 genes, had greater than 30% mitochondrial genes or had UMI counts greater than 8000 as an indicator of cell doublets. We removed genes that were detected in less than 3 cells. We normalized data using ‘SCTransform’ and used first 20 principal components with a resolution of 0.8 for clustering. We then removed computationally identified doublets from each dataset using DoubletFinder (version 2.0.3) (23). The ‘pN’ value was set to default value of 0.25 as the proportion of artificial doublets. The ‘nExP’ was set to expected doublet rate according to Chromium Single Cell 3’ version 2 reagents kit user guide (10x Genomics). These parameters were used as input to the ‘doubletFinder_v3’ function with number of principal components set to 20 to identify doublet cells.

### Batch-corrected integrated scRNA-seq analysis

Individual Seurat objects were merged and normalized using ‘SCTransform’ (64, 65). To eliminate potential batch effects, we integrated all datasets using the Harmony algorithm (version 0.1.0) (24) using patient as the grouping variable in the ‘RunHarmony’ function. Harmony reduction was used in both ‘RunUMAP’ and ‘FindNeighbors’ functions for clustering. The first 20 principal components and a resolution of 2 was used for clustering. The data from the ‘RNA’ assay was used for all further downstream analysis with other packages, gene level visualization or differential expression analysis. The data was normalized to the logarithmic scale and the effects of variation in sequencing depth were regressed out by including ‘nCount_RNA’ as a parameter in the ‘ScaleData’ function.

### Cell lineage identification and reclustering of integrated scRNA-seq data

From the batch-corrected Seurat object, cell lineages were identified based on marker gene expression. Clusters lacking marker genes but with high expression of mitochondrial or heat shock protein family genes, and those expressing markers of more than one lineage indicative of doublets were filtered from the downstream analysis (19). We performed a secondary clustering analysis of each lineage with integration across patients using Harmony and a cluster resolution of 1. Any clusters identified as belonging to another cell lineage were united with their lineage counterparts for a second clustering run. This yielded final lineage-specific re-clustering results. In integrated analysis of T0 samples, a single proliferative cluster with both B and T cells was gated for T cells based on the expression of normalized counts for *CD3D* or *CD3E* > 0.

Clusters containing T and NK cells were subjected to further cell type identification. To ensure elimination of B-T doublets, we filtered cells expressing immunoglobulin genes as described previously (19). Immunoglobulin gene expression was quantified using Seurat ‘AddModuleScore’ function and cells with expression score >0 were filtered. T and NK cell lineages were identified by reference mapping to the single-cell Tumor infiltrating immune cell atlas (21). Seurat object and metadata for the atlas was obtained from 10.5281/zenodo.4263972 and filtered for cells belonging to T and NK cell lineages. Counts from the reference atlas were normalized to the logarithmic scale and used as a reference for automated annotation per cell using SingleR (version 1.14.1) (25). Raw counts were used to annotate test datasets. Labels were predicted for each cell in the test dataset using the ‘SingleR’ function to calculate the Spearman correlation for 50 marker genes for the reference dataset identified with Wilcoxon Rank Sum test. Following automated label assignment using this method, we confirmed results by examining marker gene expression. Cell labels were then reannotated in case of misassignment in keeping with current recommended best practices (66). T helper cells were renamed as TFh-like cells. Cytotoxic and effector memory CD8 were renamed as cytotoxic CD8 T cells. Pre-exhausted and terminally exhausted cells were renamed as dysfunctional CD8 T cells. Naive-memory CD4 T cells were grouped together with naïve cells. Transitional memory cells were regrouped with TFh-like cells and Th17 were regrouped with cytotoxic CD8 T cells based on marker gene expression. We identified any dysfunctional CD8 cells misidentified as TFh-like cells based on normalized expression of *CD8A* or *CD8B* > 0. Finally proliferating cells belonged to multiple lineages and were identified using gating for lineage specific counts. Dysfunctional CD8 T cells were classified based on expression of *CD8A* or *CD8B* > 0 followed by Tregs with *FOXP3* >0 with the remainder as TFh-like cells. A final round of harmonized clustering was performed on control and individual treatment comparisons for lineage of interest. For example – CD8 T cells from ctrl and TIGIT conditions.

### Differential expression

Differential expression analysis between control and treated cells was conducted using model-based analysis of single-cell transcriptomics (MAST) (32) (version 1.18.0) on genes expressed in greater than 10 percent cells using log normalized data. The number of detected genes were recalculated after filtering. MAST hurdle model was modelled for the treatment condition and adjusted for the number of detected genes. To account for inter-patient variability, we incorporated sample as a random effect in the linear mixed model. This was implemented using the ‘zlm’ function in MAST with ‘glmer’ as the method and ‘ebayes’ set to false. A threshold of log fold change of 0.25 and false discovery rate (FDR) *p* < 0.05 was used to identify significant DE genes. In cases where we examined differences between groups of cells without modelling interpatient variability, we used the ‘FindAllMarkers’ or ‘FindMarkers’ Seurat functions for differential expression using Wilcoxon rank sum test. These instances are identified in the manuscript text. Parameters provided for these functions were as follows: genes detected in at least 25% cells and differential expression threshold of 0.25 log fold change. Significant genes were determined with *p* < 0.05 following Bonferroni correction.

### Pathway analysis

Gene sets of interest were obtained from MSigDB Human Collections (67) using package msigdbr (version 7.5.1). These included ‘BIOCARTA_NFKB_PATHWAY’ and ‘REACTOME_TCR_SIGNALING’ from curated gene sets, ‘GO_CELLULAR_RESPONSE_TO_CALCIUM_ION’ and ‘GOBP_T_CELL_ACTIVATION’ from biological process gene ontology, ‘HALLMARK_INTERFERON_GAMMA_RESPONSE’ from Hallmark gene sets. Additional gene sets for cytotoxic effector gene signature (35, 36) and CD8 dysfunction (26) were compiled from literature (**Supplemental Table 3**). We used the ‘AddModuleScore’ function in Seurat to calculate the expression of a gene set of interest in each cell using default parameters. Expression between categories was compared using unpaired t-test. Effect size was estimated using Cohen’s d measure with equal variance and Hedge’s correction implemented in rstatix (version 0.7.0).

### Differential abundance analysis

We detected cells with differential abundance between conditions using DA-seq version 1.0.0 (37). All steps from the vignette were performed on harmonized cell embeddings with a resolution of 0.01 in the function ‘getDAregions’

### TCR analysis

Filtered Cell Ranger V(D)J outputs indicative of productive TCR chains detected in high-confidence cells were used. Only the TRB chain was used in downstream analysis to define clonotype of a cell. In cases of multiple TRB sequences detected per cell, we retained the sequence with higher UMI count as described previously (19). In rare instances with ties for UMI count, we retained both sequences per cell. Expansion index in cell types using Shannon entropy was calculated with R package Startrac (version 0.1.0) (18).

### Target expression in public datasets

Single-cell Tumor infiltrating immune cell atlas from 13 cancer types (21) was used to visualize target gene expression. To evaluate expression in our previously published GC dataset (17), we reference mapped T and NK cells to the Tumor infiltrating immune cell atlas as outlined above. For visualization in both datasets, all CD4 subtypes were grouped as Naïve cells, T Helper cells were renamed as ‘CD4_TFh’, cytotoxic and dysfunctional subsets were grouped as described above.

### Multiplex Immunofluorescence

Antibodies used for multiplex immunofluorescence (mIF) staining included CD8α (C8/144B, #70306, 1:800), TIGIT (E5Y1W, #99567T, 1:800), FOXP3(D2W8E, # 98377, 1:200), GITR (D9I9D, #68014, 1:400) and Signal Stain Boost IHC Detection reagents for species specific HRP conjugated secondary antibodies (all from Cell Signaling Technology). TSA Plus Fluorescein, Cyanine 5, and Cyanine 3 kit (Akoya Biosciences) were used for tyramide signal amplification. Staining was carried out as per manufacturer’s protocol (Cell Signaling Technology). Briefly, FFPE sections were deparaffinized in Histochoice clearing agent and hydrated in a descending alcohol series. Antigen retrieval was performed with boiling 1 mM EDTA, pH 8.0 using a microwave with maintenance at a sub-boiling temperature for 15 min. Staining order, antibody concentrations and fluorophore combinations were optimized using a human tonsil section obtained from the Stanford Tissue Bank. Order of antibodies and fluorophores in one panel was GITR (Cy3), FOXP3 (Cy5) and CD8 (Fluorescein) used sequentially. Another panel comprised TIGIT (Cy3), FOXP3 (Cy5) and CD8 (Fluorescein) used sequentially. Stripping of antibodies following signal amplification was performed using boiling 10 mM Sodium Citrate, pH 6.0 in a microwave followed by maintenance at sub-boiling temperature for 10 minutes and cooling on bench top for 30 minutes. Nuclear staining was performed with 2 μg/ml DAPI (Thermo Fisher Scientific).

### RNA In situ hybridization

RNA-ISH was performed for *GZMB* using the RNAscope Multiplex Fluorescent Reagent kit v2 (ACD BioTechne) as per manufacturer’s protocol for FFPE sections. TSA Plus Cyanine 5 was used for detection. Staining pattern was confirmed in a human tonsil section. Positive and negative control probes supplied by the manufacturer were used to evaluate signal to noise ratio in the tonsil section.

### Image analysis

Images were acquired on a Zeiss Axio Imager Widefield Fluoresce Microscope (Stanford Neuroscience Microscopy Service) from two or three representative regions of interest per sample. Image analysis was performed in QuPath version 0.3.2 (68). Cell detection was performed with default parameters except minimum area was set to 5 μm2. Composite classifier was created for mIF staining in each sample using mean signal intensity thresholds per cell for each fluorophore. Steps outlined in the QuPath vignette were followed. Cells positive for both FOXP3 and CD8 were filtered (1.88-2.67% of total cells). For RNA-ISH analysis, number of spots per cell was counted using subcellular detection function in QuPath.

### Additional statistical analysis and visualization

We used the Adjusted Rand Index (ARI) to compare similarity between cluster labels and condition batch meta data label for each cell. A vector of these respective class labels was supplied to the ‘adjustedRandIndex’ function in mclust package (version 5.4.7). Additional analysis or visualization was conducted using R packages stats (version 4.1.0.), tibble (version 3.1.7), dplyr (version 0.7.6), broom (version 0.7.6), ggplot2 (version 3.3.6), ggpubr (version 0.4.0) and ComplexHeatmap (version 2.9.3) (69) in R version 4.1.0 (70). Seurat functions ‘DimPlot’ and ‘DotPlot’ were also used for visualization. Figures were additionally edited in Adobe Illustrator CS6 (version 16.0.0).

## Data availability

Sequencing data deposition is in progress under dbGAP identifier phs001818. Cell Ranger matrices will be available on https://dna-discovery.stanford.edu/research/datasets/.

## Authors’ contributions

AS and HPJ designed the study. AS acquired the data and developed the methodology. AS, XB and SMG analyzed the data. AS, XB and HPJ interpreted the data. CA, BL, CK, ASh, and GP provided translational support. AS and HPJ wrote the manuscript with input from all authors. HPJ supervised the study.

## Acknowledgements

We are grateful to all patients who participated in the study as well as their families. We thank Alison Almeda for assistance in sample collection and Colin Sim for assistance in RNA-ISH. Figures 1A, 5G and 7G were created using Biorender.com.

## SUPPLEMENTAL DATA

Supplemental Figures S1 – S2. Format: PDF

Supplemental Tables S1 – S7. Format: XLSX

