## Supplemental figures for "GITR and TIGIT immunotherapy provokes divergent multi-cellular responses in the tumor microenvironment of gastrointestinal cancers"

Hanlee P. Ji

Mailing address: CCSR 2245, 269 Campus Drive, Stanford, CA-94305, USA

### SUPPLEMENTAL FIGURE LEGENDS

**Supplemental Figure 1.** (A) Proportion of cell types detected from each baseline T0 sample. (B-E) Quantification of mIF across all samples and regions of interests. Error bars indicate standard error of mean. (F) Representative image from H&E staining of a 24 hour ctrl TSC from sample CRC_5. Scale bar = 100 μm.

**Supplemental Figure 2.** (A) Representative images of RNA-ISH staining for *GZMB* and nuclear stain DAPI in control or TIGIT treated TSCs from samples CRC_5 and GC_1_3. Scale bar = 20 μm. (B) Quantification of *GZMB* RNA-ISH from three representative regions of interest from control or TIGIT treated TSCs from respective samples with t-test *p*.

### Supplemental Figure 2

**A**

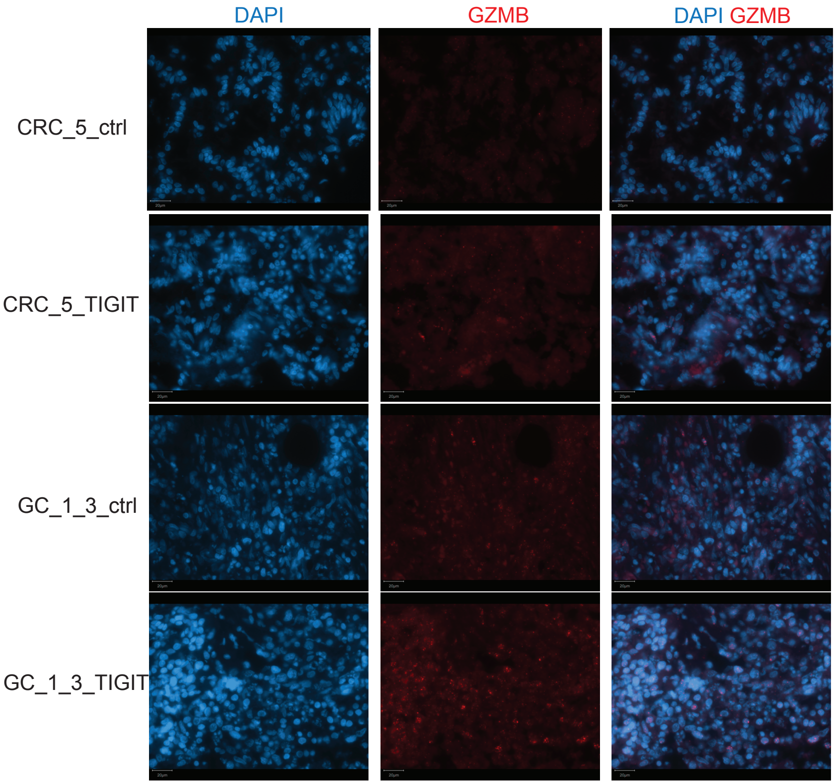


**B**

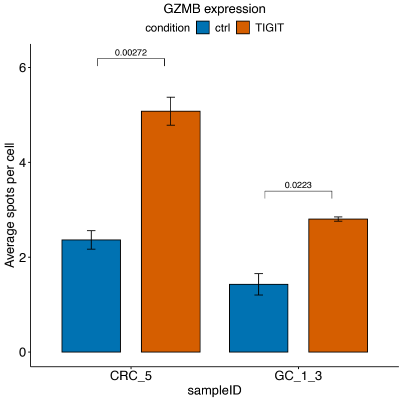

### Supplemental Figure 1

A

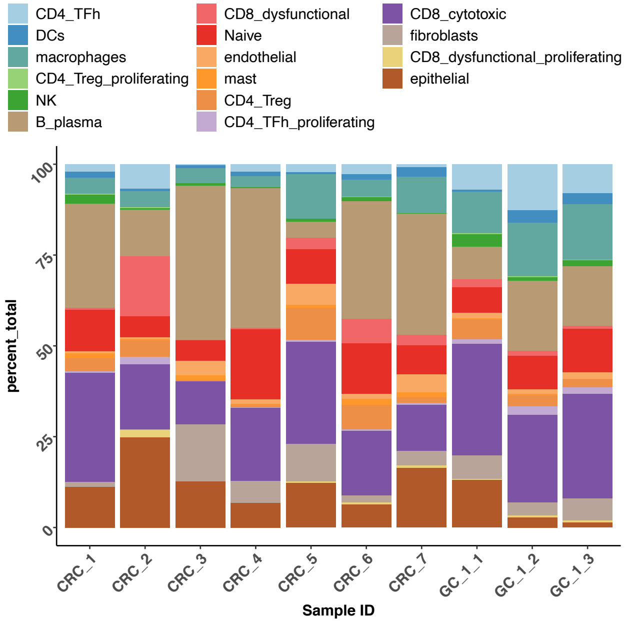


B

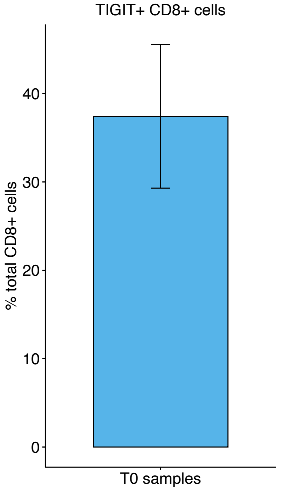


C

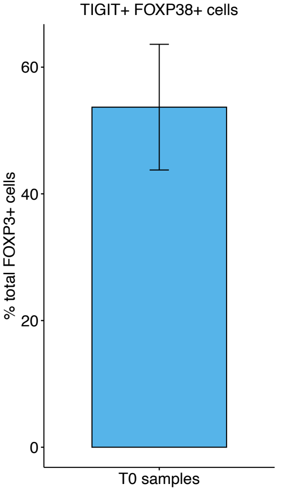


D

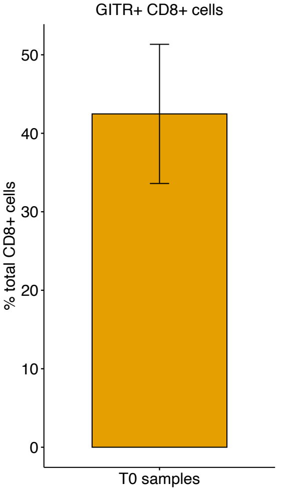


E

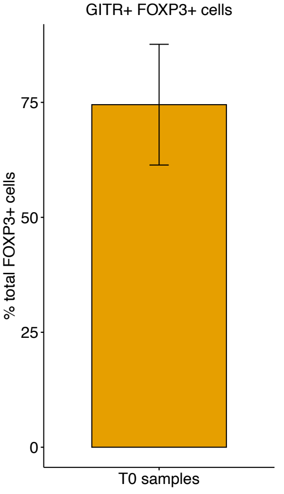


F

CRC_5 tumor slice culture

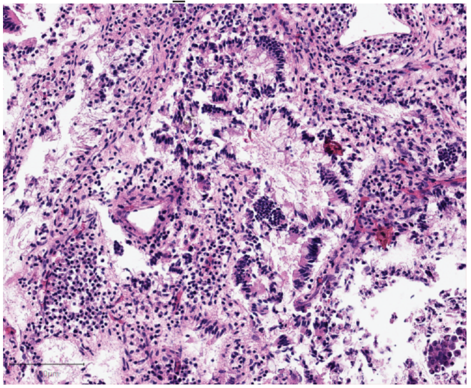
